# Comparison of sodium azide- and cooling-based immobilization methods for *Caenorhabditis elegans* chemotaxis assays

**DOI:** 10.64898/2026.09.28.755163

**Authors:** Rachel Ji, Shawndra Wibisono, Jingru Sun

## Abstract

Chemotaxis assays are widely used to quantify sensory-driven behavioral responses in *Caenorhabditis elegans*. Sodium azide is commonly used to immobilize animals at assay endpoints, although concerns have been raised that diffusion beyond the endpoint region could affect behavior before a choice is completed. We compared sodium azide-mediated immobilization with post-assay cooling using the repellent 1-octanol and attractant diacetyl. Wild-type animals showed robust avoidance of 30% 1-octanol and attraction to 0.5% diacetyl with either method, across multiple assay durations, and with or without worm food *Escherichia coli* OP50. These results support post-assay cooling as an alternative endpoint-scoring method.

## Description

Chemotaxis assays are widely used in *Caenorhabditis elegans* to investigate sensory processing, neuronal function, and behavioral responses to attractive and aversive chemical cues (1-5). In a typical chemotaxis assay, animals are placed between spatially separated test and control regions, and their distribution after a defined assay period is used to quantify attraction or avoidance (6). Because *C. elegans* remain mobile after reaching an assay endpoint, animals may subsequently leave the selected region, potentially obscuring their initial behavioral choice. Sodium azide (NaN_3_) is therefore commonly applied at assay endpoints to immobilize animals after they enter a target region and prevent redistribution before scoring (6, 7). However, a potential concern with this approach is that NaN_3_ may diffuse beyond the intended endpoint region and affect locomotion before animals have completed their behavioral response. To address this concern, we compared the conventional NaN_3_-based approach with an alternative method in which animals remained freely mobile throughout the assay and were subsequently incubated at 4°C for 15 min before scoring. Cooling at 4°C has been used to halt animal movement and preserve plate occupancy for subsequent scoring in *C. elegans* chemotaxis assays (6). We asked whether these two immobilization methods produce comparable chemotactic responses under the same experimental conditions.

We compared the two immobilization approaches using two well-characterized olfactory stimuli that elicit opposite behavioral responses in *C. elegans*: 1-octanol, an aversive odorant, and diacetyl, an attractant at low concentrations (2, 3). Synchronized wild-type N2 young adults were placed at the center of square assay plates with vehicle and test-stimulus regions positioned on opposite sides (Fig. 1A). For the sodium azide condition, NaN_3_ was applied to the endpoint regions before animals were introduced, consistent with its established use for immobilizing animals at chemotaxis assay endpoints (6). For the cooling condition, NaN_3_ was omitted and animals were allowed to move freely throughout the assay, after which the plates were incubated at 4°C for 15 min before scoring. Approximately 100 animals were tested per plate, and six independent biological replicates were analyzed for each condition.

**Figure 1.**
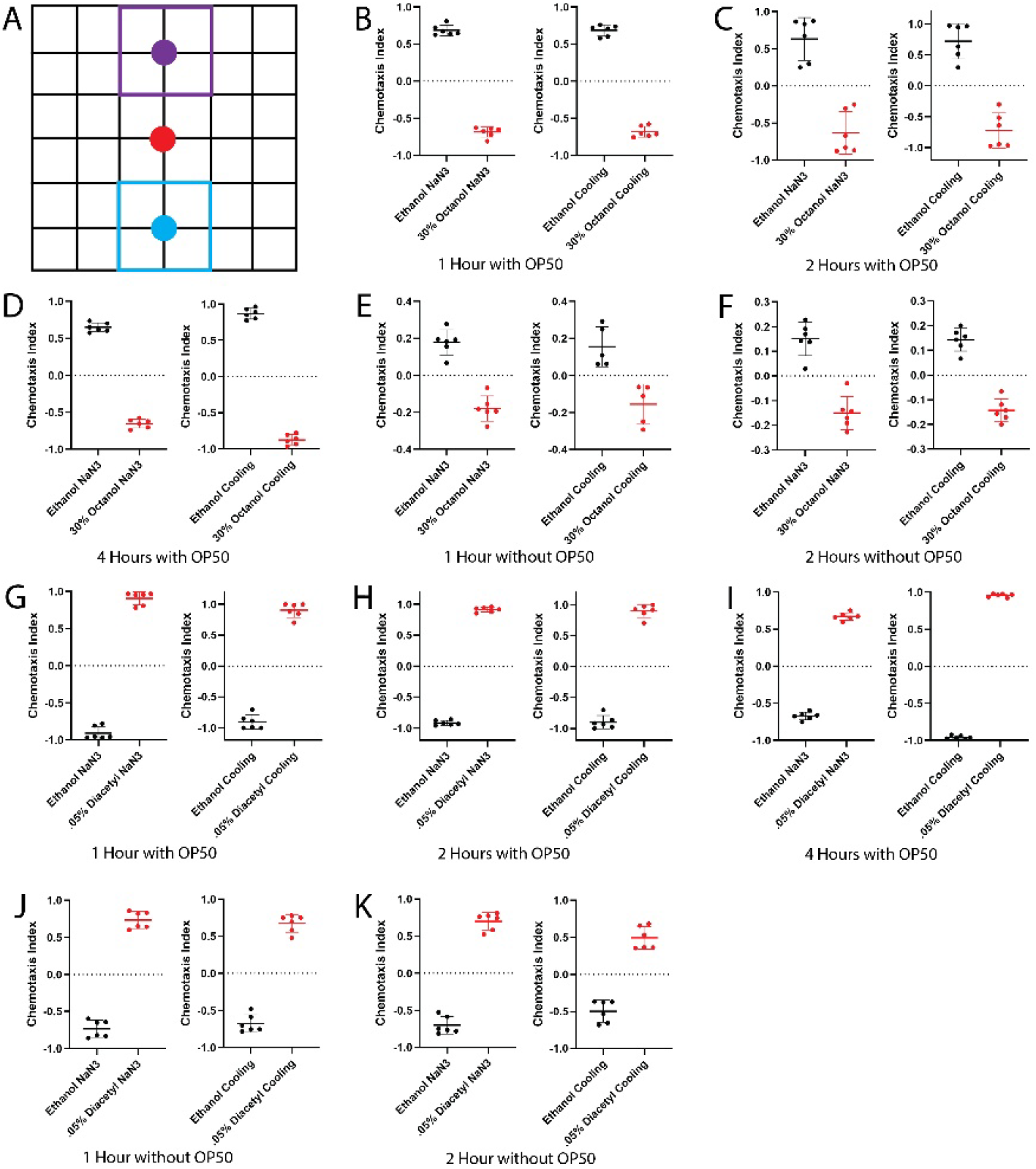
Sodium azide- and cooling-based immobilization preserve robust aversive and attractive chemotaxis responses in *C. elegans*. (A) Schematic of the chemotaxis assay. Animals were placed at the center of square assay plates (red circle), with the ethanol vehicle-associated region (blue) and test stimulus-associated region (purple) positioned at opposite ends of the plate. Animals were scored as occupying the vehicle-associated region, test stimulus-associated region, or no-decision region. For sodium azide assays, NaN_3_ was applied to the assay endpoints before animals were introduced. For cooling-based assays, NaN_3_ was omitted and plates were incubated at 4°C for 15 min before scoring. Synchronized wild-type N2 young adults were tested for behavioral responses to 30% 1-octanol (B-F) or 0.5% diacetyl (G-K). Assays were scored after 1 h (B,E,G,J), 2 h (C,F,H,K), or 4 h (D,I). Experiments were performed in the presence of OP50 (B-D,G-I) or in the absence of OP50 (E,F,J,K). Each point represents one independent biological replicate; horizontal lines and error bars indicate mean ± SD. Six biological replicates were analyzed per condition, with >80 animals scored per replicate. The choice index was calculated as (*NN*_test_ − *NN*_vehicle_)/*NN*_total_, where *NN*_test_is the number of animals in the test stimulus-associated region, *NN*_vehicle_is the number of animals in the vehicle-associated region, and *NN*_total_is the total number of animals scored on the plate, including animals in the no-decision region. Statistical significance of the behavioral response within each immobilization condition was determined using Student’s *t*-test. *P*-values were: (B) NaN_3_, *P* < 0.0001; cooling, *P* < 0.0001. (C) NaN_3_, *P* < 0.0001; cooling, *P* < 0.0001. (D) NaN_3_, *P* < 0.0001; cooling, *P* < 0.0001. (E) NaN_3_, *P* < 0.0001; cooling, *P* = 0.0019. (F) NaN_3_, *P* < 0.0001;cooling, *P* < 0.0001. (G) NaN_3_, *P* < 0.0001; cooling, *P* < 0.0001. (H) NaN_3_, *P* < 0.0001; cooling, *P* < 0.0001. (I) NaN_3_, *P* < 0.0001; cooling, *P* < 0.0001. (J) NaN_3_, *P* < 0.0001; cooling, *P* < 0.0001. (K) NaN_3_, *P* < 0.0001; cooling, *P* < 0.0001.

In assays containing the bacterial food source *E. coli* OP50, 30% 1-octanol elicited robust avoidance under both immobilization conditions. At 1 h, animals preferentially occupied the ethanol vehicle-associated region and avoided the 1-octanol-associated region whether sodium azide or cooling was used (Fig. 1B). The same behavioral pattern was maintained at 2 h (Fig. 1C) and 4 h (Fig. 1D). Thus, the characteristic aversive response to 1-octanol was preserved when sodium azide-mediated immobilization was replaced by cooling. Significant differences between occupancy of the vehicle- and 1-octanol-associated regions were observed with both immobilization approaches across these time points.

Because the presence of bacterial food could influence locomotion, residence time, or endpoint occupancy, we repeated the 1-octanol assays in the absence of OP50. Animals again preferentially occupied the vehicle-associated region and avoided the 1-octanol-associated region after 1 h and 2 h with either sodium azide or cooling (Fig. 1E,F). We did not extend the no-OP50 condition to 4 h because prolonged food deprivation is known to alter locomotor and chemotactic behavior in *C. elegans* and could therefore confound interpretation of the assay (8, 9). These findings indicate that the corresponding 1-octanol avoidance pattern was maintained in the absence of a bacterial lawn over the time window examined.

We next asked whether the concordance between the two immobilization approaches also extended to an attractive stimulus. In the presence of OP50, animals exposed to 0.5% diacetyl preferentially occupied the diacetyl-associated region relative to the vehicle-associated region at 1, 2, and 4 h (Fig. 1G-I), consistent with the known attractive effect of low-concentration diacetyl. This response was observed with both sodium azide and cooling, indicating that the direction of chemotactic behavior was preserved across immobilization methods. The same pattern was observed in the absence of OP50 at 1 h and 2 h (Fig. 1J,K). As with the 1-octanol assays, the no-OP50 diacetyl experiments were restricted to 1- and 2-h time points to minimize potential confounding effects of prolonged food deprivation.

Taken together, these experiments show that NaN_3_-mediated immobilization and post-assay cooling produced concordant qualitative behavioral outcomes for both an aversive and an attractive stimulus under the conditions tested. In each case, the direction of the chemotactic response was preserved across the two approaches. Although diffusion of sodium azide from the endpoint region is theoretically possible, the small amount used in our assay (10 µL of 30 µM NaN_3_) did not produce an apparent alteration in chemotactic behavior relative to the cooling method. Cooling nevertheless allows animals to remain freely mobile throughout the assay without exposure to a chemical paralytic and therefore provides a practical alternative when potential NaN_3_ diffusion is a concern.

These findings do not establish that cooling and sodium azide are interchangeable across all *C. elegans* chemotaxis paradigms. The present study examined wild-type young adults, 30% 1-octanol, 0.5% diacetyl, and assay durations of up to 4 h in the presence of OP50 and up to 2 h in its absence. Subtle behavioral preferences, weaker sensory phenotypes, different stimulus concentrations, developmental stages, assay geometries, or shorter-duration responses may be more sensitive to the immobilization procedure. Nevertheless, the consistent direction of behavioral responses across the two stimuli, multiple assay durations, and both food conditions supports post-assay cooling as a practical alternative to sodium azide for endpoint scoring of robust *C. elegans* chemotaxis responses.

## Methods

### *C. elegans* culture and synchronization

Wild-type *C. elegans* N2 animals were maintained on SB plates seeded with *Escherichia coli* OP50. Animals were synchronized by timed egg laying. Twenty well-fed adult animals were transferred to OP50-seeded SB plates and allowed to lay eggs for 45 min at 25°C, after which the adults were removed. Progeny were subsequently maintained at 20°C for 65 h until they reach the young adult stage.

### Chemotaxis assay

Chemotaxis assays were performed on square SB plates marked with a 6 × 6 grid to facilitate scoring. For assays containing a bacterial food source, two 10-µL drops of freshly prepared *Escherichia coli* OP50 suspension were placed on opposite sides of each plate and allowed to dry. Three 2-µL drops of either ethanol vehicle or test stimulus were then placed adjacent to the corresponding endpoint region such that animals encountered the stimulus before reaching the bacterial lawn. For aversive chemotaxis assays, 30% 1-octanol (MilliporeSigma, Cat. No. 112615) prepared in ethanol was used as the test stimulus. For attractive chemotaxis assays, 0.5% diacetyl (MilliporeSigma, Cat. No. 11038; 2,3-butanedione) was used. Parallel assays were performed in the absence of OP50 to assess whether the presence of bacterial food influenced the behavioral response.

For sodium azide-mediated immobilization assays, 10 µL of 30 µM NaN_3_ was applied to the endpoint regions and allowed to dry before animals were introduced. For cooling-based assays, NaN_3_ was omitted.

Synchronized young adult animals were collected in M9 buffer and washed three times by centrifugation at 3000 rpm for 3 min, with removal of the supernatant after each wash.

Approximately 100 animals were transferred to the center of each assay plate. Individual plates were placed in airtight containers and maintained at 20°C for the duration of the assay. Animals were scored after 1, 2, or 4 h, as indicated. For assays performed in the absence of OP50, only the 1- and 2-h time points were analyzed. For cooling-based assays, plates were incubated at 4°C for 15 min before scoring to reduce locomotion and preserve animal positions.

Animals were classified as occupying the vehicle-associated region, the test stimulus-associated region, or the no-decision region. Behavioral preference for each plate was quantified using a choice index calculated as:

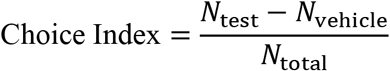

where *NN*_test_is the number of animals in the test stimulus-associated region, *NN*_vehicle_is the number of animals in the vehicle-associated region, and *NN*_total_is the total number of animals scored on the plate, including animals in the no-decision region. Because diacetyl is an attractant for *C. elegans*, positive choice-index values indicate attraction toward the diacetyl-associated region, whereas values near zero indicate little or no preference. In contrast, because 1-octanol is a repellent, avoidance is reflected by negative choice-index values, indicating preferential occupancy of the vehicle-associated region. Thus, the sign of the choice index reflects the direction of the behavioral response to the test stimulus.

## Statistical analysis

Statistical analyses were performed using GraphPad Prism version 11 (GraphPad Software, USA). Each assay plate was considered one independent biological replicate. Six independent biological replicates were analyzed for each condition, with more than 80 animals scored per replicate. For each immobilization method, behavioral responses were analyzed within each experimental condition using Student’s *t*-test, as specified in the Figure 1 legend. Statistical significance was defined as *P* < 0.05, and exact *P* values are reported in the figure legend. No direct statistical comparison was made between the sodium azide and cooling immobilization methods. Accordingly, the two approaches were evaluated on the basis of whether they produced concordant qualitative chemotactic responses under the conditions tested rather than on formal statistical equivalence.

## Acknowledgements

We thank Dr. Phillip Wibisono for technical assistance. We also thank the *Caenorhabditis* Genetics Center (CGC), which is funded by the NIH Office of Research Infrastructure Programs (P40 OD010440), for providing the *C. elegans* strain used in this study. This work was supported by NIH (R35GM124678 to J.S.). The funder had no role in study design, data collection and interpretation, or the decision to submit the work for publication.

## Author contributions

R.J. and S.W. designed the study, performed the experiments, and analyzed the data. J.S. conceived and supervised the project, contributed to experimental design and data interpretation, and wrote the manuscript.

